# Tracing the origin of non-brittle rachis alleles in wheat

**DOI:** 10.64898/2026.02.26.708176

**Authors:** Emile Cavalet-Giorsa, Thomas Wicker, Simon G. Krattinger

## Abstract

The evolution of non-brittle rachises, controlled by the *TtBTR1* genes, was a key step during wheat domestication. Here, using *k*-mer-based approaches applied to a large diversity panel, we refine previous estimates of the geographical and temporal origins of the three known *Ttbtr1* loss-of-function alleles and show that they emerged in distinct wild emmer subpopulations. For this, we generated a chromosome-scale dika wheat assembly carrying the recently discovered *Ttbtr1-Ab* allele. Our analyses reveal that the *Ttbtr1-A* alleles reside on an introgression from the southern *judaicum* wild emmer population into northern wild emmer wheat, providing an explanation for the long-standing debate about the *Ttbtr1-A* origin. We further demonstrate that the *Ttbtr1-Aa* and *Ttbtr1-B* alleles are already present in wild emmer wheat, with evidence indicating that they arose prior to the advent of agriculture. Together, these findings support a model in which key domestication genes in domesticated crops were selected and combined from standing genetic variation in wild relatives.

## Introduction

The transition to agriculture in the Near East marked one of humankind’s most profound sociocultural transformations, laying the foundation for modern civilizations. This shift was closely tied to the domestication of plants and animals. In cereals, a key target of domestication was rachis brittleness^1-4^. While the wild progenitors of cereals shed grains at maturity through brittle internodes, cultivation favored spikes that remained intact for harvesting. Key genes contributing to rachis brittleness in *Triticeae* were first discovered in barley, where loss-of-function mutations in either the *BTR1* or *BTR2* gene confer the non-brittle phenotype^3,5^.

In wheat, non-brittleness is similarly caused by loss-of-function mutations in *BTR1*. Three *Ttbtr1* alleles have been identified in domesticated wheats of the emmer lineage, which includes both durum wheat (*Triticum turgidum* ssp. *durum*) and bread wheat (*T. aestivum*), the two economically most important wheat species today. Initially, only two *Ttbtr1* alleles were known, *Ttbtr1-Aa* on chromosome 3A and *Ttbtr1-B* on chromosome 3B. The combination of *Ttbtr1-Aa* and *Ttbtr1-B* results in non-brittleness, leading to the view that the emergence of this trait in the emmer wheat lineage was monophyletic. The *Ttbtr1-Aa* allele carries a two-base-pair deletion, whereas *Ttbtr1-B* harbors a ∼4 kb insertion^2,4^. Both mutations disrupt the *TtBTR1* coding sequence. More recently, a second *Ttbtr1-A* allele, *Ttbtr1-Ab*, was discovered. It carries a 5,029 bp retrotransposon insertion and has been found in tetraploid *T. turgidum* ssp. *carthlicum* (dika wheat) as well as in some domesticated tetraploid wheat accessions from Ethiopia, revealing two independent origins of the *Ttbtr1-A* allele^6^. The exact evolutionary origins of *Ttbtr1* and domesticated wheat remain contested^7-9^, a debate further complicated by the mosaic haplotype composition of cereal genomes shaped by extensive gene flow among populations^10-13^.

## Results and Discussion

To investigate the origin and distribution of the recently discovered *Ttbtr1-Ab* allele^6^, as well as the evolution of non-brittleness, we first generated a chromosome-scale assembly of the *Ttbtr1-Ab*-carrying *T. turgidum* ssp. *carthlicum* accession CWI 22960 by combining PacBio HiFi reads with Hi-C. The assembly spanned 10.88 Gb with a contig N50 of 42.6 Mb (Supplementary Table 1). In this accession, *Ttbtr1-Ab* carried the 5,029 bp retrotransposon insertion at position 459 bp, with two identical 248 bp long terminal repeats (LTRs). Next, we performed an allelic diversity analysis at *Ttbtr1-A* and *Ttbtr1-B* using a *k*-mer database derived from whole-genome sequencing data of 2,130 domesticated and 463 wild wheat accessions (Supplementary Table 2, 3), including both tetraploids and hexaploids. In total, 98.6% of the domesticated accessions carried the 2 bp deletion characteristic for *Ttbtr1-Aa*. In contrast, only 30 (1.4%) domesticated wheat accessions harbored the recently identified *Ttbtr1-Ab* allele. The 30 accessions represent tetraploid wheats, 13 of which are classified as *T. turgidum* ssp. *carthlicum* (dika wheat), 13 domesticated Ethiopian tetraploids, two *T. turgidum* ssp. *durum*, one *T. turgidum* ssp. *polonicum*, and one *T. turgidum* ssp. *turanicum* (Supplementary Table 2).

No additional loss-of-function allele was found for *Ttbtr1-B*. Among the 463 wild emmer wheat accessions, nine carried the *Ttbtr1-Aa* allele (*Ttbtr1-Aa* /*TtBtr1-B*), and seven wild emmer wheat accessions had the *Ttbtr1-B* allele (*TtBtr1-A* /*Ttbtr1-B*) (Supplementary Table 3). Five wild emmer wheat accessions caried both *Ttbtr1-Aa* and *Ttbtr1-B*, which can be due to accession misclassification or gene flow from domesticated wheat. No wild emmer wheat lines were found carrying the newly identified *Ttbtr1-Ab* allele.

Wild emmer wheat can be grouped into three main clades, (i) a northeastern population comprising accessions collected from present-day southern Anatolia, Iran, and Iraq (EST_WEW), (ii) a Southern Levant population (LEV_WEW), and (iii) race *judaicum* (JUD_WEW) found around the Sea of Galilee^4,6,14^. A *k*-mer-based phylogeny across the 463 wild emmer wheat accessions and 30 tetraploid domesticated wheat accessions recovered these three major wild emmer wheat clades. The LEV_WEW population further split into two subgroups (LEV_WEW-1 and LEV_WEW-2) along the north-south gradient, while the EST_WEW population resolved into five subgroups (EST_WEW-1 to EST_WEW-5), highly associated with the geographical provenance of the accessions from west to east (Fig. 1a, b; Supplementary Figs. 1-3; Supplementary Table 3).

**Figure 1.**
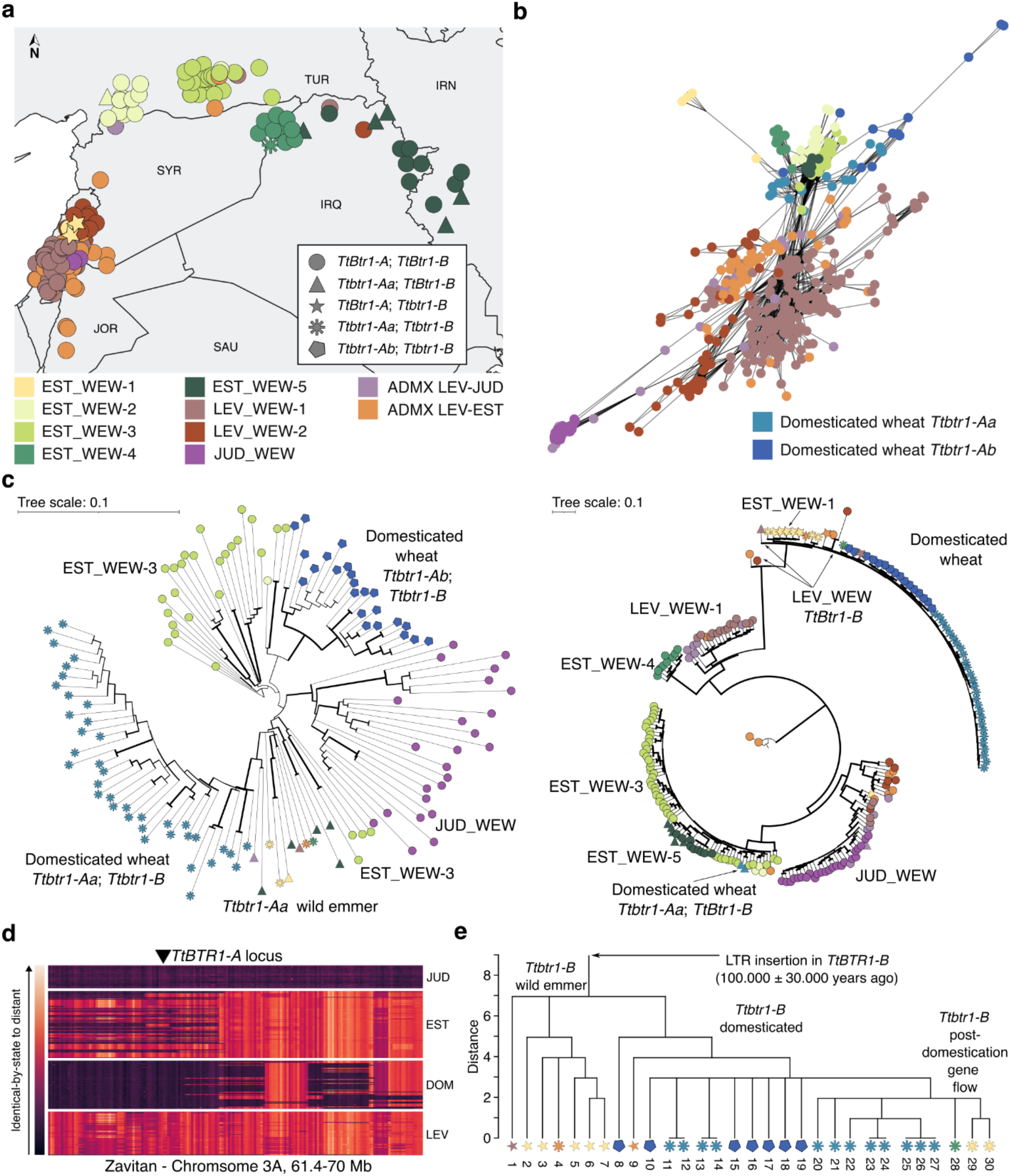
Origin of non-brittle rachis in wheat. **a**, Map showing the collection sites and the wild emmer wheat accessions analyzed in this study. Only accessions with known collection sites are shown. IRQ, Iraq; IRN, Islamic Republic of Iran; JOR, Hashemite Kingdom of Jordan; SAU, Kingdom of Saudi Arabia; SYR, Syrian Arab Republic; TUR, Türkiye. Jittering is applied for map readability. Colors for the different subpopulations and symbols for the *TtBtr1* allele combinations will be used throughout the figure. **b**, *k*-mer-based network representation of the wild emmer wheat population structure. Accessions with a normalized distance closer than 0.61 are connected. **c**, *k*-mer-based phylogenetic trees across the 300-kb genomic segments surrounding the *TtBTR1-A* (left) and the *TtBTR1-B* loci (right). For *TtBTR1-A* (left) only samples showing identity-by-state across the whole locus are shown to increase the resolution. For the *TtBTR1-B* locus (right), only a subset of accessions from distinct populations is reported for the readability of the tree. Bootstrap support values are represented by node thickness. **d**, Genome identity heatmap across chromosome 3A (position 61.4-70 Mb). The wild emmer wheat Zavitan, belonging to the *judaicum* race, was used as a reference. Dark regions indicate identity-by-state to Zavitan, while red color indicates genomic segments that are distant from Zavitan. The *Ttbtr1-Aa* and *Ttbtr1-Ab* alleles are located on a *judaicum* introgression that is widespread across the northern wild emmer wheat group (EST_WEW) but absent in the Southern Levant wild emmer wheat group (LEV_WEW). **e**, Phylogenetic tree obtained comparing the SNPs presents on the long terminal repeats (LTRs) of the transposable element inserted in the *Ttbtr1-B* allele. Samples can be identified by the numbers reported in the panel and in Supplementary Tables 2 and 3.

To further refine the phylogeny of the *TtBtr1* loci, we selected non-recombining 300-kb genomic segments surrounding *TtBTR1-A* and *TtBTR1-B* and constructed two phylogenetic trees for the complete diversity panel based on identity-by-state scores, using the assembly of Chinese Spring^15^ (hexaploid wheat, *Ttbtr1-Aa* /*Ttbtr1-B*) as a reference (Fig. 1c, Supplementary Fig. 4, 5). For *Ttbtr1-A*, the domesticated wheat accessions carrying *Ttbtr1-Aa* and *Ttbtr1-Ab* clustered into two distinct branches of the phylogenetic tree (Fig. 1c). Most of the wild emmer wheat accessions with *Ttbtr1-Aa* belong to the northern subpopulation EST_WEW-5 from Iran and Iraq, whereas the closest wild emmer wheat relatives with the wild-type *TtBtr1-A* allele belong to the northern subpopulation EST_WEW-3 from southern Anatolia, for both *Ttbtr1-Aa* and *Ttbtr1-Ab* (Fig. 1c). This broader northern distribution region has previously been recognized as the most likely origin of *Ttbtr1-Aa*, although its precise geographic origin and subpopulation could not be determined^7-9^. Notably, we found that *Ttbtr1-Aa* and *Ttbtr1-Ab* reside within a genomic introgression from the *judaicum* subpopulation from Southern Levant. This introgression has become widespread across northern wild emmer wheats (Fig. 1d), accounting for the difficulty in pinpointing the exact origin of *Ttbtr1-A*, but is mostly absent in the Southern Levant wild emmer wheat. The widespread distribution of the *judaicum* introgression in EST_WEW is consistent with the previously reported low genetic diversity at the *Ttbtr1-A* locus^6^. The longest version of the *judaicum* introgression was found in CWI 22960 and two other *T. turgidum* ssp. *carthlicum* accessions, spanning around 140 Mb.

Consistent with previous reports, the Levant region represents the most likely origin of *Ttbtr1-B*. Five of the seven wild emmer wheat accessions carrying *Ttbtr1-B* fall within the EST_WEW-1 subpopulation. While most of the accessions belonging to EST_WEW-1 have an unknown collection site, the accessions with known geographical location were all from Lebanon. The genetically closest wild emmer wheat accessions with the wild-type *TtBtr1-B* allele belong to the Southern Levant subpopulations. The five genetically closest wild emmer wheat accessions with wild-type *TtBtr1-B* were collected from Lebanon (3 accessions), Syria, and Israel (Fig. 1c). Taken together, these phylogenetic patterns point to the northern margin of the Southern Levant region as the most probable origin of *Ttbtr1-B*.

The occurrence of *Ttbtr1-Aa* and *Ttbtr1-B* in wild emmer wheat may reflect either post-domestication gene flow from domesticated wheat or standing genetic variation that predates agriculture, analogous to the *teosinte glume architecture* (*tga1*) allele in maize^16^ and the *btr1* allele in barley^10^. To estimate the timing of *Ttbtr1-B* emergence, we dated the associated retrotransposon insertion by evaluating SNPs between its two long terminal repeats^17^. The previously reported ∼4kb insertion in *Ttbtr1-B* was an underestimation, resulting from the limitations of short-read-based genome assemblies^4^. *Ttbtr1-B* sequences from 13 long-read-based domesticated wheat assemblies (Supplementary Table 4) revealed a 11.97 kb retrotransposon insertion with two long terminal repeats of approximately 3,894 bp (with minor size differences caused by homopolymers). In addition, we mapped raw reads from the seven wild emmer wheat accessions carrying *Ttbtr1-B* (*TtBtr1-Aa* /*Ttbtr1-B*), four wild emmer wheat accessions carrying both *Ttbtr1-Aa* and *Ttbtr1-B*, and six domesticated tetraploid wheat accessions to the Chinese Spring reference genome^15^. SNPs were then called across the LTRs. Phylogenetic analyses revealed that some *Ttbtr1-B*-carrying wild emmer wheat accessions clustered with domesticated wheat, consistent with post-domestication gene flow, while other wild emmer wheat accessions formed a distinct clade defined by private SNPs unique to this group (Fig. 1e). Among the latter are TA11213 and GT004, two wild emmer wheat accessions from Lebanon belonging to subpopulation EST_WEW-1. LTR dating based on the 13 chromosome-scale assemblies indicated a transposon insertion age of ∼100,000 ± 30,000 years (Supplementary Table 5). While molecular dating relies on assumptions about mutation rates^17^, even our most recent estimate (49,000 ± 22,000 years) places the origin of *Ttbtr1-B* around 27,000 years ago, well before the beginning of agriculture. A pre-domestication origin of *Ttbtr-1B* is further supported by the SNP-based phylogenetic analysis, which showed that some LTR-specific SNPs occur exclusively in wild emmer wheat and are absent from domesticated wheat. In addition, this timing is consistent with the finding of domestic-type wild emmer wheat spikes at the Ohalo II archaeological site before the advent of agriculture^18^. The long terminal repeats of the retrotransposon inserted in *Ttbtr1-Ab* were too short for dating. We thus used a SNP-based dating approach^10^, estimating the emergence of the *Ttbtr1-Aa* and *Ttbtr1-Ab* mutations approximately 38,000 and 33,600 years ago.

Together, our findings support a model in which key mutations for rachis brittleness arose in different wild emmer wheat subpopulations and were maintained in natural or early human-managed settings. Wild emmer wheat accessions carrying a single non-brittle rachis allele showed brittle/semi-brittle rachises (Fig. 2). Plants with semi-brittle rachises have been found in stands of wild emmer wheat, and it has been reported that environmental factors such as humidity and temperature likely influence spike maturation and shattering^19^. The presence of a single non-brittle rachis allele in wild emmer wheat therefore likely has no strong fitness effect and spikes will eventually shatter. The combination of *Ttbtr1-A* and *Ttbtr1-B* alleles through hybridization, resulting in fully domesticated wheat with non-brittle rachis, likely occurred multiple times, giving rise to distinct wheat subspecies.

**Figure 2.**
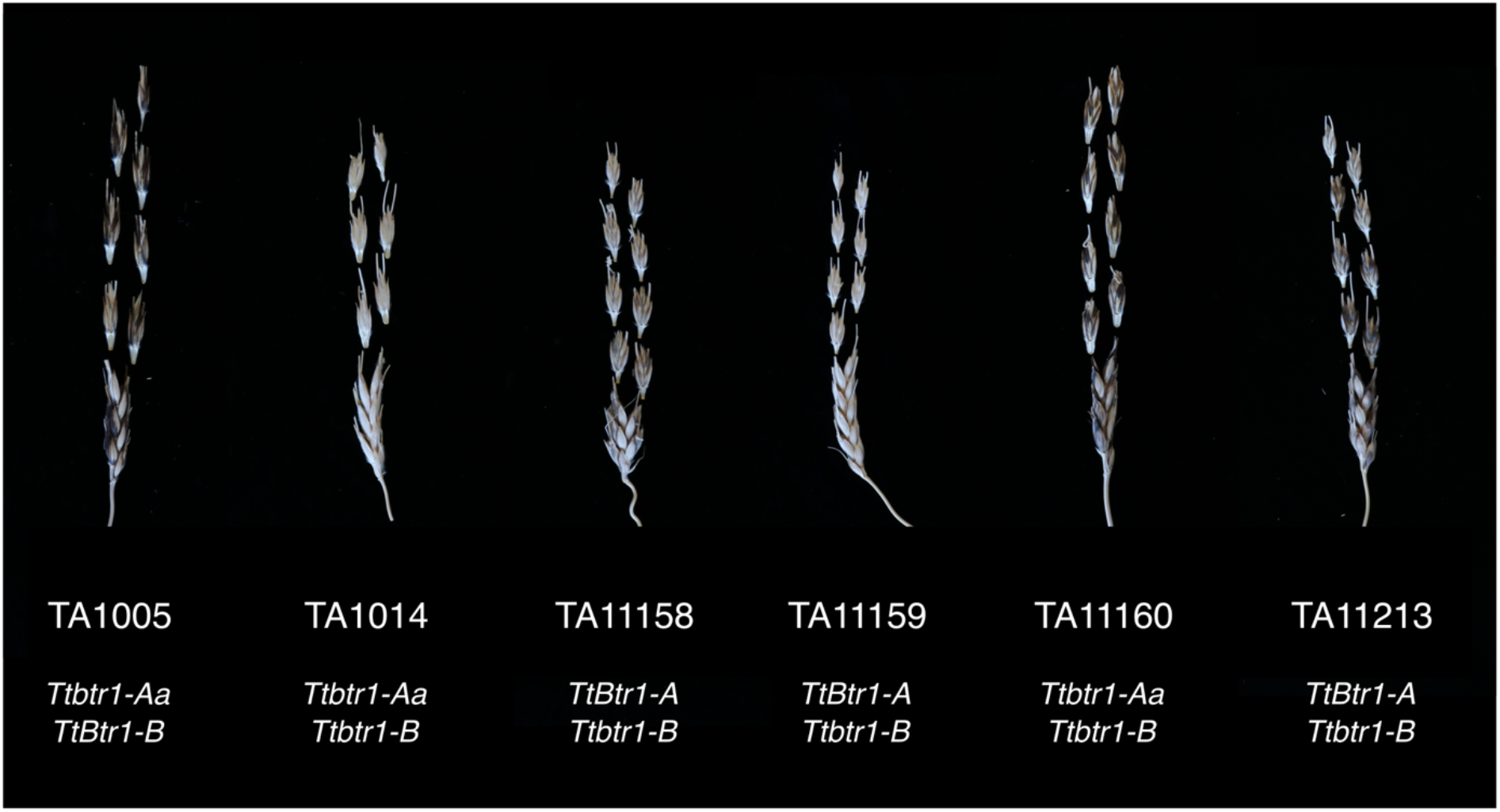
Phenotypic representation for wild emmer wheat spikes carrying *Ttbtr1-Aa, Ttbtr1-B* or both at the same time. All spikes show a semi-brittle phenotype comparable to the one of domesticated emmer.

## Methods

### *T. turgidum* ssp. *carthlicum* (CWI 22960) assembly

High molecular weight DNA was extracted from dark-treated 2-week-old seedlings using the Qiagen Genomic Tip kit. The output of four PacBio Revio SMRT cells (in total 364.62 Gb, 33.5-fold coverage, 23.99 million reads, 15,211 bp average read length) was assembled with hifiasm (v. 0.19.8)^20^ with default settings. The 799 million Illumina short reads generated by the sequencing of a Hi-C library were first mapped to the CWI 22960 contig-level assembly with BWA-MEM using the Arima Genomic mapping pipeline’s default settings (https://github.com/ArimaGenomics/mapping_pipeline). The resulting file was then processed with YaHS (v. 1.1)^21^. A few rounds of manual curations were performed with the help of Juicebox (v. 2.15)^22^ and CHROMEISTER^23^. PacBio Revio sequencing was performed in the KAUST Bioscience Core Lab. Hi-C library preparation and sequencing was done by CNRGV as a service.

### Whole-genome sequencing

Genomic DNA from the 25 wheat accessions was extracted from one or two young leaves of a single plant following the CTAB protocol described by Abrouk *et al*.^24^. PCR-free library preparation and sequencing at 12-fold coverage were performed as a service by Psomagen Inc. (Rockville, Maryland, United States).

### Whole-genome sequencing data collection

Publicly available wheat sequencing data were retrieved from NCBI from the following project numbers: PRJNA1007489, PRJNA759292, PRJNA628827, PRJNA476679, PRJNA310175, PRJNA688544, PRJNA1070409, PRJNA1188632, PRJNA439156, PRJNA476679, PRJNA596843, PRJNA597250, PRJNA663409, PRJNA669381, PRJNA670578, PRJNA694980, PRJNA714281, PRJNA722149, PRJNA729723, PRJNA744310, PRJNA745496, PRJNA759292, PRJNA771357, PRJNA790490, PRJNA820989, PRJNA877303, PRJNA900700, PRJNA918327, PRJNA956839, PRJNA986484, PRJNA986532; from EBI-ENA from the following project numbers: PRJEB61424, PRJEB22687, PRJEB44721, PRJEB45541, PRJEB48529, PRJEB49351; and from the Genome Sequence Archive in the BIG Data Center from the following project numbers: PRJCA004228, PRJCA004273, PRJCA005979, PRJCA009783, PRJCA019508, PRJCA019636, PRJCA021345.

### *k*-mer counting

Raw Illumina reads were cleaned with Trimmomatic (v. 0.39)^25^ with the following settings: LEADING:3 TRAILING:3 SLIDINGWINDOW:4:25 MINLEN:75. 31-mers were counted with KMC3 (v. 3.1.2)^26^.

### *k*-mer-based phylogeny

The pairwise intersections between *k*-mer sets representing different accessions were computed with the FastIBS KDBIntersect function (https://github.com/githubcbrc/FastIBS). To account for different coverages, which influences the total number of *k*-mers in a set, we applied a ‘reduction factor’ to each comparison value depending on the number of total *k*-mers present in each of the two datasets. To compute the ‘reduction factor’, a Random Forest Regressor was trained on the surface obtained with the intersections of *k*-mer sets, simulating different coverages for two datasets (CRR061704 and SRR11670754). For CRR061704, 19 datasets corresponding to 4.5-fold to 16-fold coverage were obtained, while for SRR11670754, 28 datasets corresponding to 4.5-fold to 26-fold coverage were obtained (Supplementary Fig. 6). This process was executed with a Python (v. 3.8.8) script with the package scikit-learn (v. 1.2.2; https://github.com/emilecg/btr_analysis).

PCA and Hierarchical clustering were executed with a Python (v. 3.8.8) script with the package scikit-learn (v. 1.2.2).

The results obtained from the ‘reduction factor’ subtraction were then normalized to have values in a 0 to 1 range. To achieve this, we applied the following formula, where X is the value of the normalized comparison, Min is the minimum value obtained from all the normalized comparisons, and Max is the maximum value obtained from all the normalized comparisons:

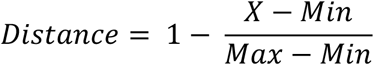

The network from the distance matrix was built using the Python package NetworkX (v3.4.2)^27^.

### Fastibsmapper – allelic diversity analysis

In order to perform the allele diversity analysis, we first used *TtBtr1-A* and *TtBtr1-B* gene sequences from Zavitan^4^ as query to search the genome of CWI 86942 using BLAST (v. 2.16.0)^28^ and extracted with Samtools (v. 1.16.1)^29^.

The gene sequences were used as reference for a FastIBS fastibsmapper run against all the *k*-mer sets generated in this study. The resulting files were stacked in two tables, plotted with a Python script (available at https://github.com/emilecg/btr_analysis) and were visually inspected to determine allelic diversity across the accessions.

### FastIBS

FastIBS (https://github.com/githubcbrc/FastIBS) was used to identify wild emmer wheat accessions carrying a non-recombinant, identical-by-state haplotype block surrounding *TtBtr1-Aa* and *TtBtr1-B* using Chinese Spring as a reference, and *TtBtr1-Ab* using the CWI 22960 assembly as a reference. Based on the FastIBS results, plotted as heatmaps, we were able to identify the non-recombining wild emmer wheat accessions for the 300-kb genomic segment surrounding the three loci. The three 300-kb loci from Chinese Spring and CWI 22960 where extracted from their respective assemblies using Samtools (v. 1.16.1)^29^ to be used as references for the local phylogeny analyses.

### Local phylogeny for *TtBtr1-A* and *TtBtr1-B*

Three 300-kb and one 30-kb segments surrounding *TtBtr1-A* and *TtBtr1-b* were used as reference for a FastIBS fastibsmapper run using all the *k*-mer sets generated in this study. The generation of the phylogenetic tree from the stacked output of fastibsmapper required four distinct steps. In the first step, all the reported drops in *k*-mer coverage with a maximum depth of 25 and a width of 5 bases were stored as a separated file. We defined a drop in *k*-mer coverage following the described criteria as a ‘valley’. This file was inspected to discard valleys that occurred in a single accession, all the other valleys were retained and saved in an output file. Two valleys were considered the same if their start and end positions occurred within 5 bases. This file was then processed in the third step: the whole list of valleys was organized in a presence absence matrix. The matrix was then converted manually into a PHYLIP format and given to IQ-TREE (v. 3.0.1)^30^ as an input. A first tree was produced to assess the general topology. After that, two more informative trees were produced with a subset including all wild emmer wheat accessions and a subset of the domesticated wheat accessions considering the very high level of similarity among this second group. The model with the best fit was selected by IQ-TREE (GTR2+FO+R4) to construct the tree. Bootstrap support values were calculated from 1,000 replicates. The tree plot was made with the online tool iTOL (v. 7.2.2)^31^. The trees were further pruned for tree readability. The three first steps were implemented with three different Python scripts available at https://github.com/emilecg/btr_analysis.

### Time estimation of the *Ttbtr1-B* retrotransposon insertion

Clustal Omega (v. 1.2.4)^32^ was used to align the LTR sequences extracted from 13 PacBio-based genomes (Supplementary Table 4). Raw reads from eleven wild emmer wheat accessions and six domesticated tetraploid accessions were mapped to the Chinese Spring assembly^15^ using BWA-MEM (v. 0.7.17)^33^ and SNPs were called over the two LTRs using bcftools mpileup (v. 1.16)^29^ with default settings. A manual inspection of the mapping results was performed to select true SNPs. The time estimation of the retrotransposon’s insertion into *Ttbtr1-B* was performed according to Wicker *et al*. 2022^17^.

### Time estimation of the *Ttbtr1-A* loss-of-function mutation events

This estimate is based on the work of Guo *et al*. 2025^10^ in barley. Whole-genome sequencing reads from 61 accessions (35 wild emmer wheat accession and 26 domesticated wheat accessions) carrying a non-recombinant identical-by-state haplotype in the *Ttbtr1-A* locus were mapped to the Zavitan genome using Minimap2 (v. 2.24)^34^ using default short-read settings. Bcftools (v. 1.16)^29^ was used to call SNPs retaining the ones with the following quality settings: -q 20 -Q 20. A filtering step was then added to retain only biallelic SNPs. Homozygous SNPs showing a depth lower than 2 and higher than 50 were set to missing. The heterozygous SNPs were set to missing if the depths of both alleles were not greater or equal than three. We then used Beagle (v. 5.4)^35^ to phase the SNP matrix to be used as an input for GEVA (v. 1)^36^. We added two artificial SNPs: one in the position of the 2 bp deletion causing the *Ttbtr1-Aa* allele and another in the position of the retrotransposon insertion causing the *Ttbtr1-Ab* allele. We then used GEVA to infer the age of the two haplotypes surrounding the two causal variants using default settings and the mutation rate determined in *Brachypodium distachyon* (6.13 × 10^-9^). Finally, we doubled the ages estimated by GEVA considering the highly homozygous nature of the wheat genome as it was done for barley by Guo *et al*.^10^. We reported the average of the results of 20 independent runs of the haplotype ages determined by the molecular clock model.

## Data availability

The Illumina whole-genome sequencing data generated in this study, the Hi-C reads generated from CWI 22960, and the CWI 22960 assembly are available at ENA under BioProject number PRJEB101210. The PacBio HiFi reads are available at ENA under BioProject number PRJEB101209.

### Code availability

All the custom python scripts used are available at https://github.com/emilecg/btr_analysis.

## Acknowledgements

We thank Natalia Arango López for the DNA extractions. We thank Yael Lev-Mirom, Assaf Distelfeld, and Curtis Pozniak for critical feedback. This publication is based upon work supported by KAUST award ORFS-CRG12-2024-6474.

## Author Contributions

S.G.K. and E.C.-G. conceived the experiments and wrote the manuscript. E.C.-G assembled the *T. turgidum* ssp. *carthlicum* genome, built the phylogeny and analyzed the *TtBtr1* alleles. T.W. performed the dating of the *Ttbtr1-B* loss-of-function event.

## References

1. Pourkheirandish, M. et al. On the origin of the non-brittle rachis trait of domesticated einkorn wheat. Frontiers in Plant Science 8, 2031 (2018).

2. Nave, M. et al. Wheat domestication in light of haplotype analyses of the Brittle rachis 1 genes (BTR1-A and BTR1-B). Plant Science 285, 193–199 (2019).

3. Pourkheirandish, M. et al. Evolution of the grain dispersal system in barley. Cell 162, 527–539 (2015).

4. Avni, R. et al. Wild emmer genome architecture and diversity elucidate wheat evolution and domestication. Science 357, 93–97 (2017).

5. Civáñ, P. & Brown, T.A. A novel mutation conferring the nonbrittle phenotype of cultivated barley. New Phytologist 214, 468–472 (2017).

6. Lev-Mirom, Y. et al. Desert cave grains uncover ancient tetraploid wheat dispersion routes. Nature Plants in press(2026).

7. Lev-Mirom, Y. & Distelfeld, A. Where was wheat domesticated? Nature Plants 9, 1201–1202 (2023).

8. Zhao, X.B. et al. Population genomics unravels the Holocene history of bread wheat and its relatives. Nature Plants 9, 403–419 (2023).

9. Zhao, X., Guo, Y. & Lu, F. Reply to: Where was wheat domesticated? Nature Plants 9, 1203–1206 (2023).

10. Guo, Y. et al. A haplotype-based evolutionary history of barley domestication. Nature (2025).

11. Cavalet-Giorsa, E. et al. Origin and evolution of the bread wheat D genome. Nature 633, 848–855 (2024).

12. He, F. et al. Exome sequencing highlights the role of wild-relative introgression in shaping the adaptive landscape of the wheat genome. Nature Genetics 51, 896–904 (2019).

13. Wang, Z.H. et al. Dispersed emergence and protracted domestication of polyploid wheat uncovered by mosaic ancestral haploblock inference. Nature Communications 13, 3891 (2022).

14. Adhikari, L. et al. Dissecting the population structure, diversity and genetic architecture of disease resistance in wild emmer wheat (Triticum turgidum subsp. dicoccoides). Research Square, 10.21203/rs.3.rs-4909521/v1 (2024).

15. Liu, S.C. et al. A telomere-to-telomere genome assembly coupled with multi-omic data provides insights into the evolution of hexaploid bread wheat. Nature Genetics 57, 1008–1020 (2025).

16. Fairbanks, R.A. & Ross-Ibarra, J. An ancient origin of the naked grains of maize. Proc Natl Acad Sci U S A 122, e2503748122 (2025).

17. Wicker, T. et al. Transposable element populations shed light on the evolutionary history of wheat and the complex co-evolution of autonomous and non-autonomous retrotransposons. Advanced Genetics 3, 2100022 (2022).

18. Snir, A. et al. The origin of cultivation and proto-weeds, long before neolithic farming. PLOS ONE 10, e0131422 (2015).

19. Peleg, Z., Abbo, S. & Gopher, A. When half is more than the whole: Wheat domestication syndrome reconsidered. Evolutionary Applications 15, 2002–2009 (2022).

20. Cheng, H.Y., Concepcion, G.T., Feng, X.W., Zhang, H.W. & Li, H. Haplotype-resolved de novo assembly using phased assembly graphs with hifiasm. Nature Methods 18, 170–175 (2021).

21. Zhou, C., McCarthy, S.A. & Durbin, R. YaHS: yet another Hi-C scaffolding tool. Bioinformatics 39, btac808 (2022).

22. Durand, N.C. et al. Juicebox provides a visualization system for Hi-C contact maps with unlimited zoom. Cell Systems 3, 99–101 (2016).

23. Pérez-Wohlfeil, E., Diaz-del-Pino, S. & Trelles, O. Ultra-fast genome comparison for large-scale genomic experiments. Scientific Reports 9, 10274 (2019).

24. Abrouk, M. et al. Fonio millet genome unlocks African orphan crop diversity for agriculture in a changing climate. Nature Communications 11, 4488 (2020).

25. Bolger, A.M., Lohse, M. & Usadel, B. Trimmomatic: a flexible trimmer for Illumina sequence data. Bioinformatics 30, 2114–2120 (2014).

26. Kokot, M., Długosz, M. & Deorowicz, S. KMC 3: counting and manipulating k-mer statistics. Bioinformatics 33, 2759–2761 (2017).

27. Hagberg, A.A., Schult, D.A. & Swart, P.J. Exploring network structure, dynamics, and function using NetworkX. in Proceedings of the 7th Python in Science Conference (SciPy2008) (eds Varoquaux, G., Vaught, T. & Millman, J.) 11–15 (Pasadena, CA USA, 2008).

28. Camacho, C. et al. BLAST+: architecture and applications. BMC Bioinformatics 10, 421 (2009).

29. Danecek, P. et al. Twelve years of SAMtools and BCFtools. Gigascience 10, giab008 (2021).

30. Wong, T.K.F. et al. IQ-TREE 3: phylogenomic inference software using complex evolutionary models. EcoEvoRxiv, 10.32942/X2P62N (2025).

31. Letunic, I. & Bork, P. Interactive Tree of Life (iTOL) v6: recent updates to the phylogenetic tree display and annotation tool. Nucleic Acids Research 52, 78–82 (2024).

32. Sievers, F. et al. Fast, scalable generation of high-quality protein multiple sequence alignments using Clustal Omega. Molecular Systems Biology 7, 539 (2011).

33. Li, H. Aligning sequence reads, clone sequences and assembly contigs with BWA-MEM. arXiv preprint arXiv:1303.3997 (2013).

34. Li, H. Minimap2: pairwise alignment for nucleotide sequences. Bioinformatics 34, 3094–3100 (2018).

35. Browning, B.L., Tian, X., Zhou, Y. & Browning, S.R. Fast two-stage phasing of large-scale sequence data. The American Journal of Human Genetics 108, 1880–1890 (2021).

36. Albers, P.K. & McVean, G. Dating genomic variants and shared ancestry in population-scale sequencing data. PLOS Biology 18, e3000586 (2020).

