## Supplementary materials for "Tracing the origin of non-brittle rachis alleles in wheat"

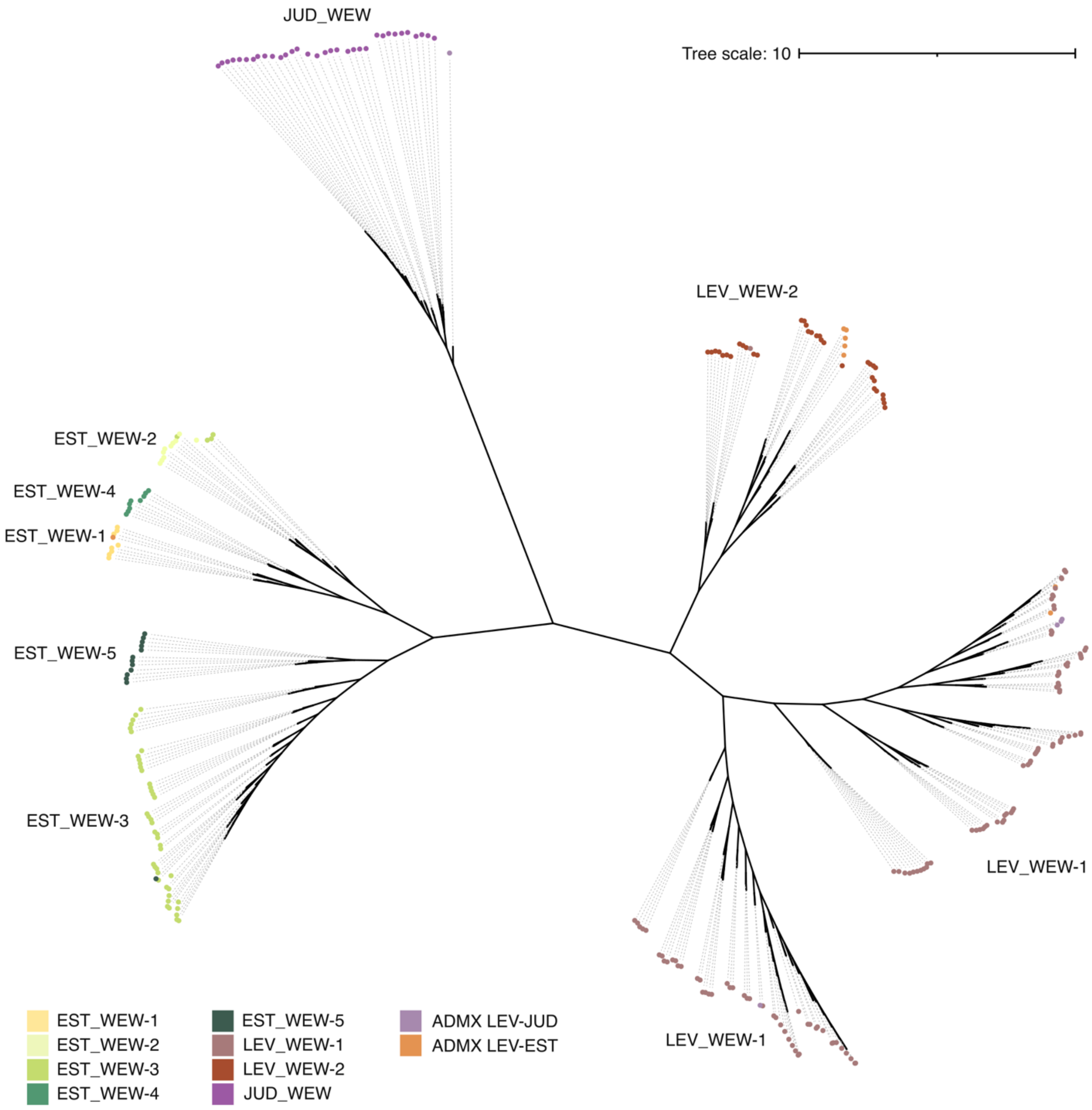


**Supplementary Fig. 1.** Hierarchical Ward's linkage tree built from the distance matrix based on *k*-mers. Only the subset of wild emmer wheat accessions included in Adhikari *et al*. 2025 are shown. The recovered tree topology is concordant with the SNP-based phylogeny presented in Adhikari *et al*. 2025.

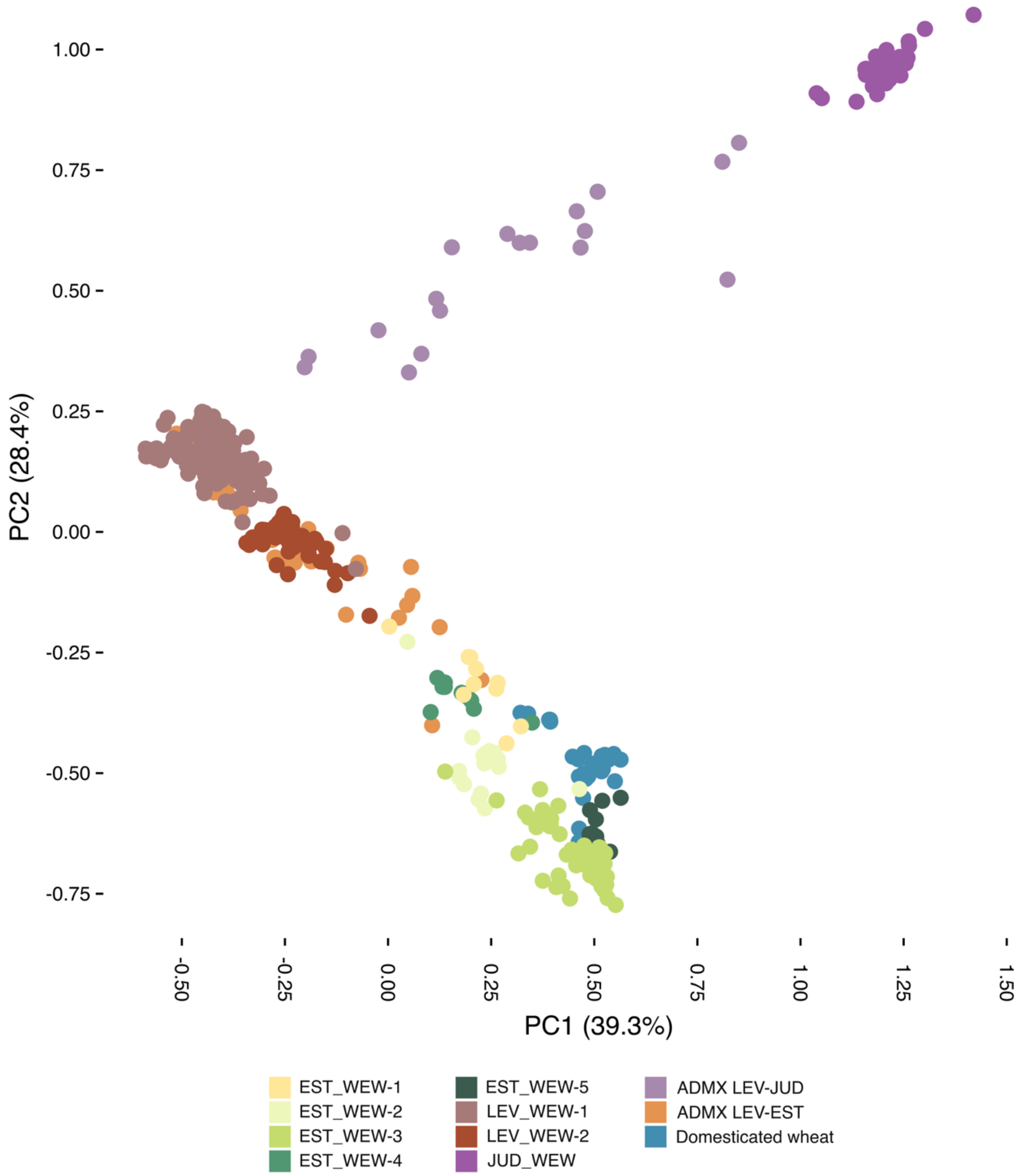


**Supplementary Fig. 2.** PCA representation of the *k*-mer distance matrix.

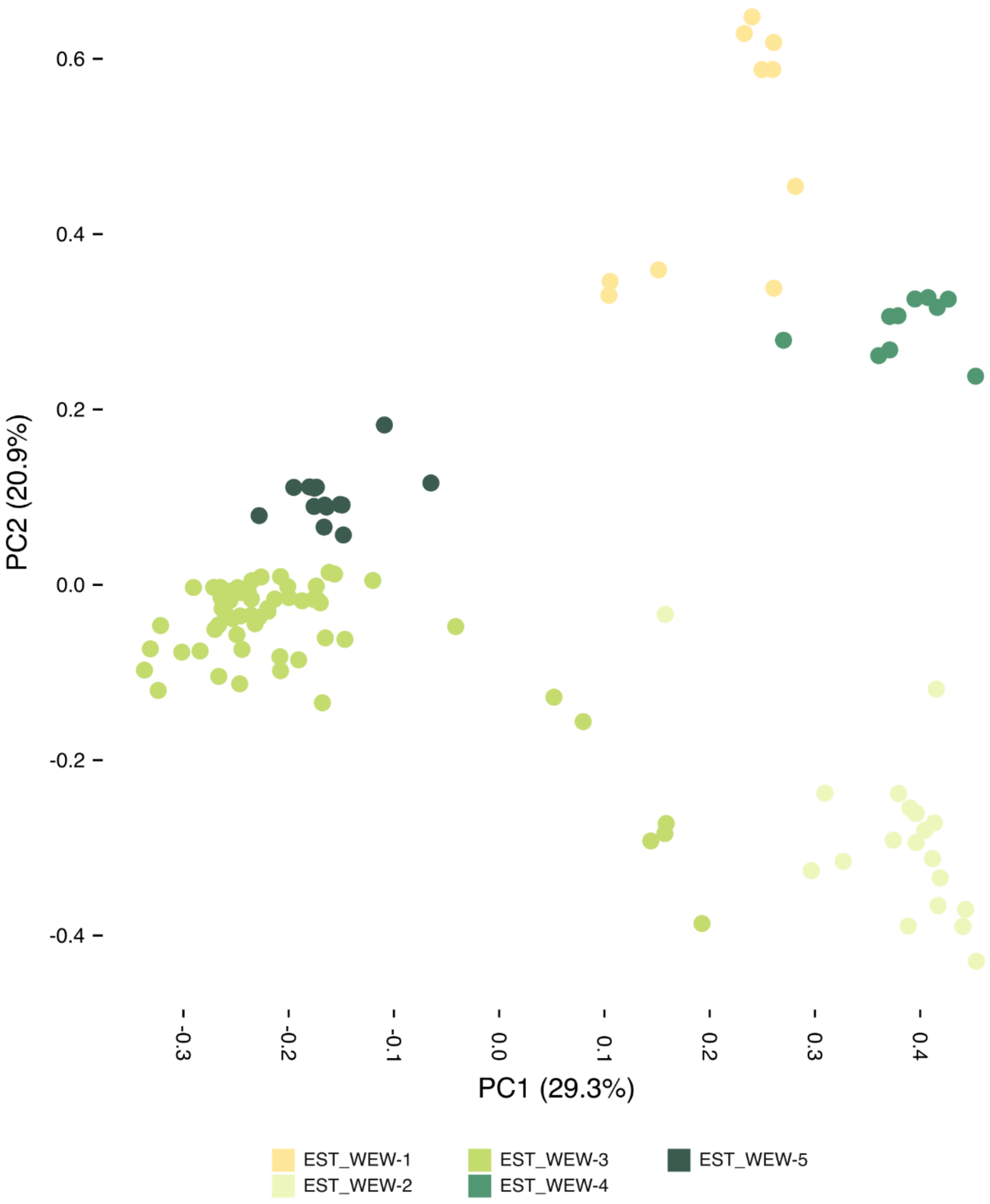


**Supplementary Fig. 3.** PCA representation of the *k*-mer distance matrix. Only the EST_WEW are plotted to show the distribution of the five subgroups.

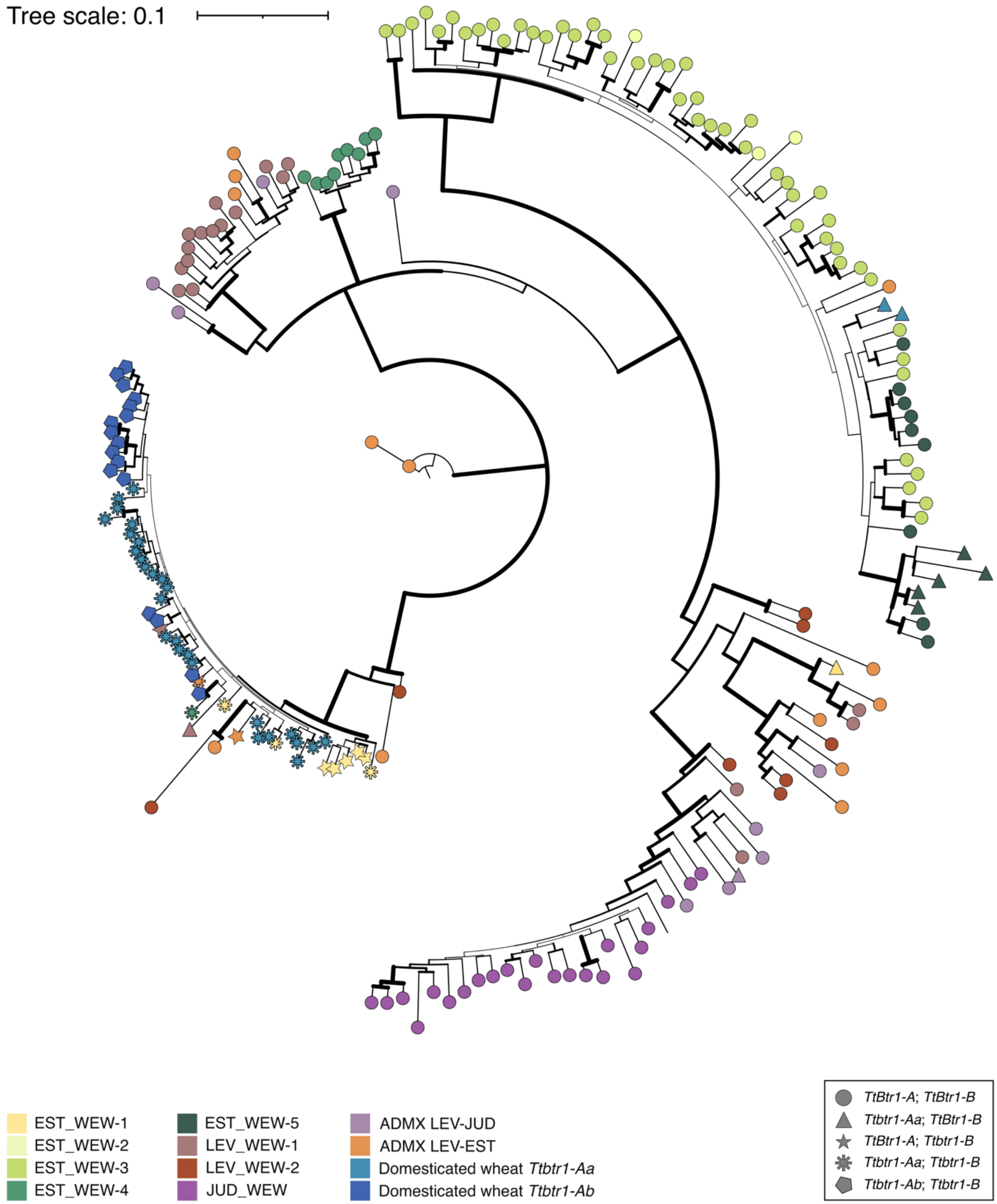


**Supplementary Fig. 4.** *k*-mer-based phylogenetic tree across a 30-kb genomic segment surrounding *TtBTR1-B*. Only a subset of accessions is reported for the readability of the tree. Bootstrap support values are represented by node thickness. The plot shows consistency to the one built over the 300-kb region surrounding *TtBTR1-B*.

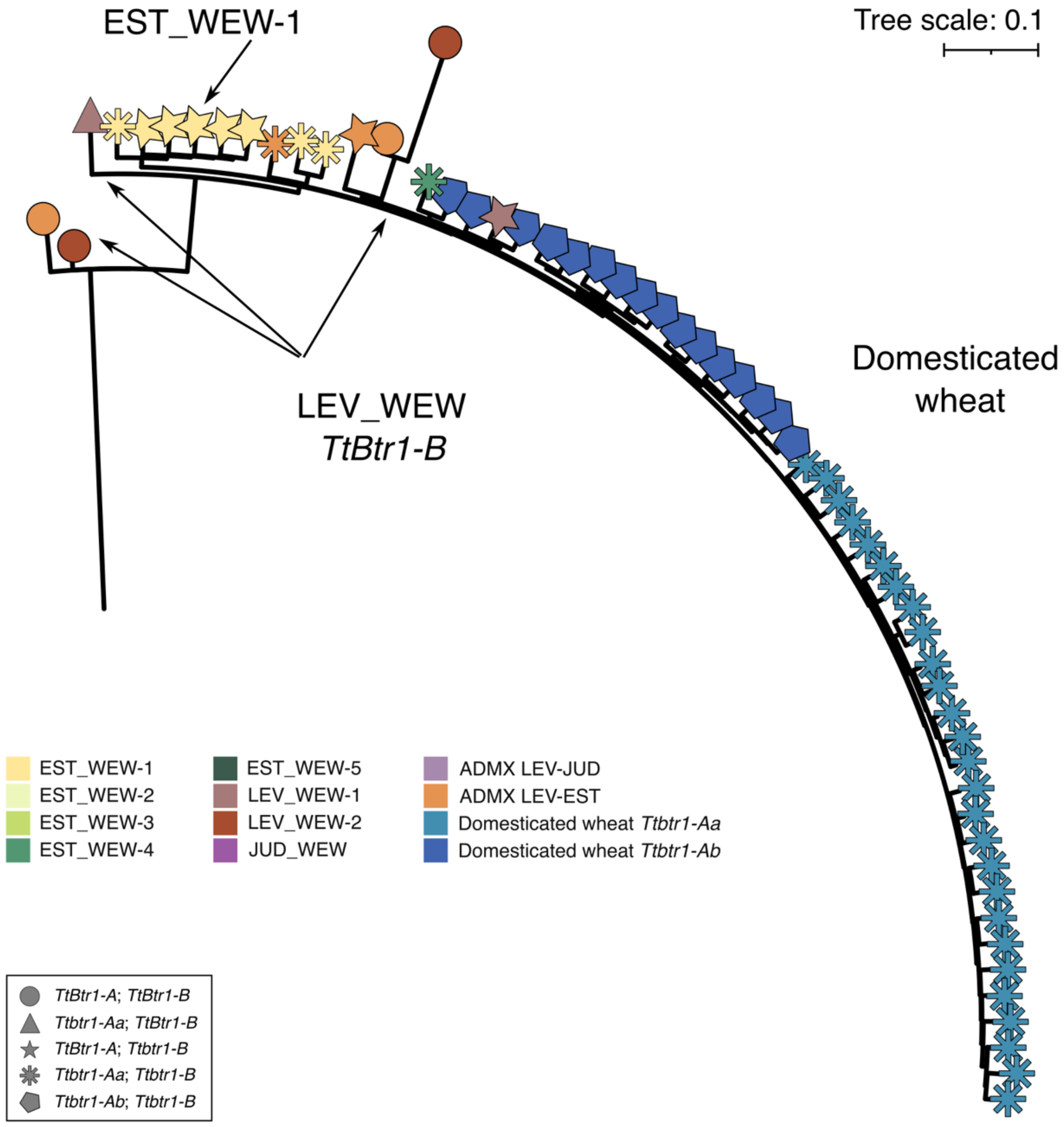


**Supplementary Fig. 5.** *k*-mer-based phylogenetic trees across a 300-kb genomic segment surrounding *TtBTR1-B* same as shown in Fig. 1c. Only an arm is reported to increase the readability of the tree.

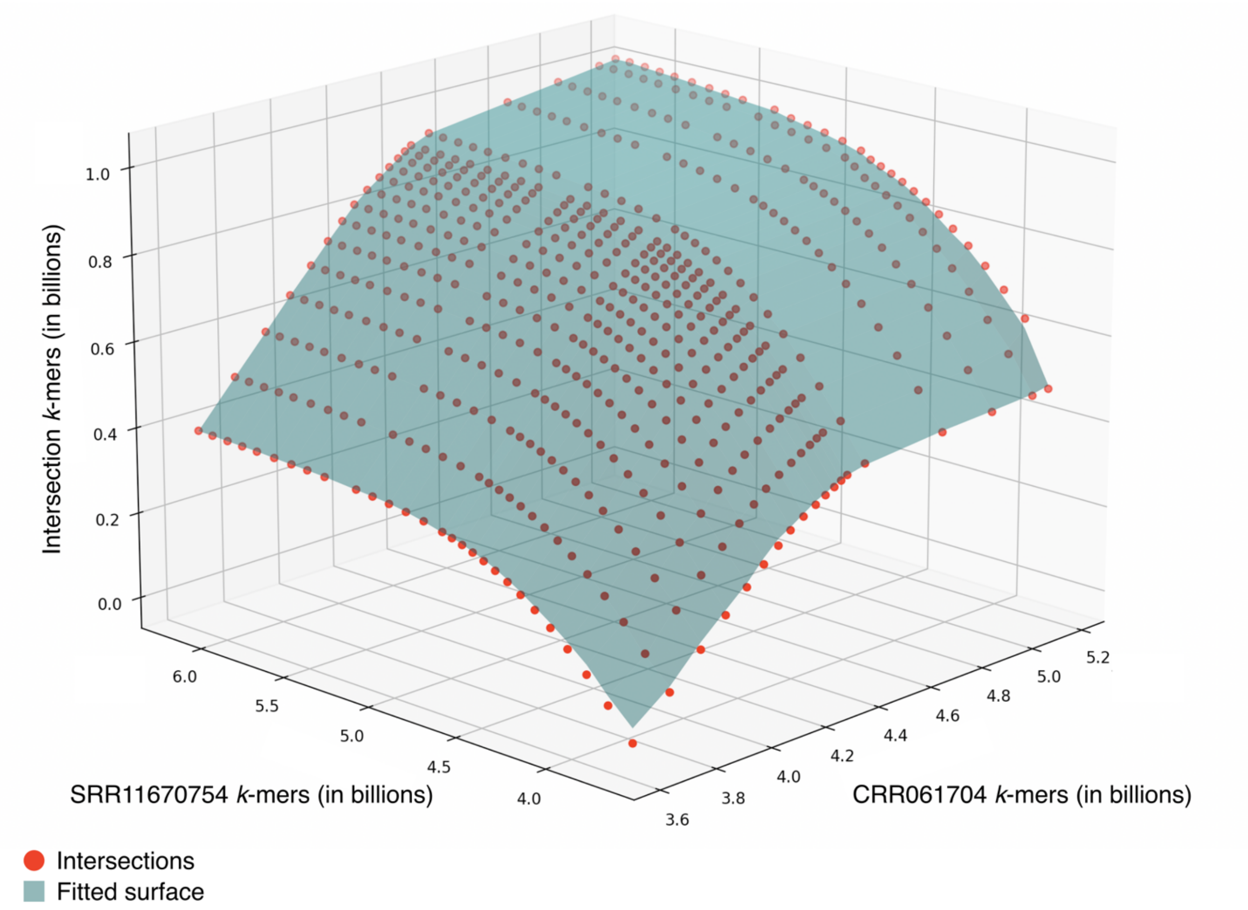


**Supplementary Fig. 6.** Representation of the intersections (red dots) between reduced *k*-mer sets to find the appropriate reduction factor for whole *k*-mers database comparisons. The fitted surface is represented in blue.
